# A Methodology for Vertical Translation Between Molecular and Organismal Level in Biological Feedback Loops

**DOI:** 10.1101/2021.09.20.461028

**Authors:** J. W. Dietrich

## Abstract

Feedback loops are among the primary network motifs in living organisms, ensuring survival via homeostatic control of key metabolites and physical properties. However, from a scientific perspective, their characterization is unsatisfactory since the usual modelling methodology is incompatible with the physiological and biochemical basis of metabolic networks. Therefore, any “vertical translation”, i.e. the study of the correspondence between molecular and organismal levels of causality, is difficult and in most cases impossible.

As a viable solution, we demonstrate an alternative modelling platform for biological feedback loops that is based on key biochemical principles, including mass action law, enzyme kinetics, binding of mediators to transporters and receptors, and basic pharmacological properties. Subsequently, we show how this framework can be used for translating from molecular to systems-level behaviour.

Basic elements of the proposed modelling platform include Michaelis-Menten kinetics defining nonlinear dependence of the output y(t) on an input signal x(t) with the Hill-Langmuir equation *y*(*t*) = *G* * *x*(*t*)^*n*^ / (*D* + *x*(*t*)^*n*^), non-competitive inhibition for linking stimulatory and inhibitory inputs with *y*(*t*) = *G* + *x*_1_(*t*) / ((*D* + *x*_1_(*t*) * (1 + *x*_2_(t) / *K*_*I*_)) and processing structures for distribution and elimination.

Depending on the structure of the feedback loop, its equifinal (steady-state) behaviour can be solved in form of polynomials, with a quadratic equation for the simplest case with one feedback loop and a Hill exponent of 1, and higher-grade polynomials for additional feedback loops and/or integer Hill exponents > 1. As a companion to the analytical solution, a flexible class library (CyberUnits) facilitates computer simulations for studying the transitional behaviour of the feedback loop.

Unlike other modelling strategies in biocybernetics and systems biology, this platform allows for straightforward translation from the statistical properties of single molecules on a “microscopic” level to the behaviour of the whole feedback loop on an organismal “macroscopic” level. An example is the Michaelis constant D, which is equivalent to (*k*_–1_ + *k*_2_) / *k*_1_, where *k*_1_, *k*_–1_ and *k*_2_ denote the rate constants for the association and dissociation of the enzyme-substrate or receptor-hormone complex, respectively. From the perspective of a single molecule the rate constants represent the probability (per unit time) that the corresponding reaction will happen in the subsequent time interval. Therefore 1/*k* represents the mean lifetime of the complex. Very similar considerations apply to the other described constants of the feedback loop.

In summary, this modelling technique renders the translation from a molecular level to a systems perspective possible. In addition to providing new insights into the physiology of biological feedback loops, it may be a valuable tool for multiple disciplines of biomedical research, including drug design, molecular genetics and investigations on the effects of endocrine disruptors.

## INTRODUCTION

In life sciences, the function of organisms uses to be described on several and distinct levels of causality, ranging from the behaviour of single elementary particles to the performance of whole living creatures and even social interactions. While approaches remaining on one level only are able to successfully provide physiological insights and while they are usable for practical purposes, including clinical reasoning and epidemiological decision making, the translation between the levels of causality is difficult and, in most cases, virtually impossible [1]. A methodology bridging this gap is highly needed, but previous approaches were unsatisfactory [2].

An example of this dilemma that is both typical and significant is feedback control systems. Feedback loops are fundamental network motifs and information processing structures in living organisms, providing means for homeostatic control of vital parameters including the concentration of key metabolites and important physical properties. Unfortunately, the usual methodology for the mathematical description of feedback loops as linear time-invariant (LTI) systems is not readily compatible with biochemical insights and theories [2]. It is, therefore, impossible to draw conclusions from physicochemical processes to the corresponding response of the organism and vice versa.

Here, we suggest an alternative modelling platform for systems biology that is able to establish compatibility between the distinct levels of causal description, thereby providing a framework for vertical translation between molecular and organismal levels of interaction. Additionally, we demonstrate how this framework can be exploited to translate between different levels of causality.

## METHODS

Based on (but not restricted to) requirements of endocrinology and metabolism we developed a metamodel, containing elements that can be mapped to empirically testable biochemical and physiological data [3, 4]. This modelling platform has been successfully applied to different hormonal feedback loops, including thyroid homeostasis, the hypothalamus-pituitary-adrenal axis and insulin-glucose homeostasis [3, 5, 6]. Its main elements include (1) coupling the concentration of a substance (e.g. a hormone or metabolite) to its elimination and distribution and modelling this process as analog signal memory with intrinsic adjustment (ASIA element) [7], (2) saturation kinetics based on the Michaelis-Menten-Hill formalism representing enzymatic processes, receptor kinetics and other forms of signal transduction mechanisms [8, 9] and (3) non-competitive inhibition for the description of negative feedback [8, 9].

ASIA elements describe the steady-state concentration of a substance with

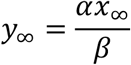

as a function of an input signal *x*_∞_, the inverse of the volume of distribution (*α*) and a clearance exponent (rate constant) *β* that depends with

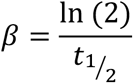

on the half-life of the respective substance [7]. The transitional behaviour is described with

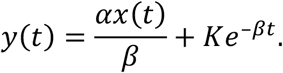

Michaelis-Menten (MiMe) kinetics describe the nonlinear dependence of an output signal *y*(*t*) on an input signal *x*(*t*) with

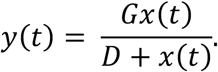

They are a well-characterised universal formalism for modelling enzymatic processes and receptor-mediated signal-transduction mechanisms.

The combined effect of stimulating (*x*_1_) and inhibiting (*x*_2_) input signals can be modelled as non-competitive divisive inhibition (NoCoDI) process with

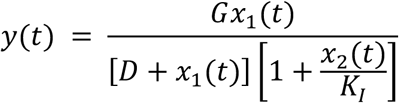

with *K*_*I*_ being the dissociation constant of *x*_2_ [3, 4].

Putting these elements together delivers a closed formalism (MiMe-NoCoDI loop) describing a nonlinear feedback control system based solely on elements that are mappable to basic biochemical or pharmacological properties (Fig. 1). The constant parameters of the feedback loop belong to different classes of physical properties: *G*_1_, *G*_2_ and *G*_3_ are extensive properties, i.e. they depend on the size of the organism, whereas *D*_2_ is an intensive property, independent of the body mass or size.

**Figure 1.**
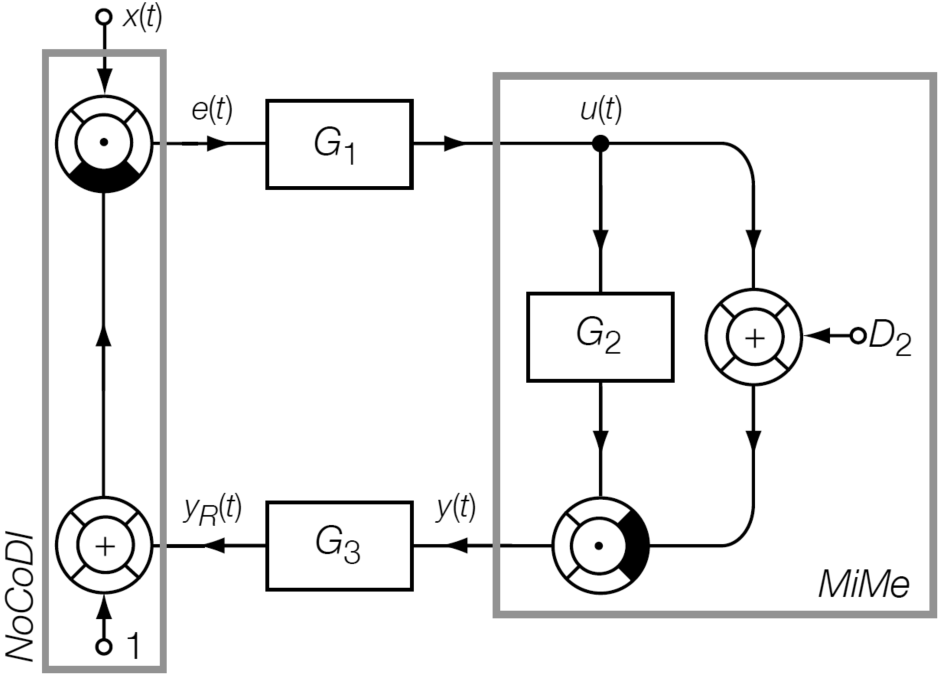
The smallest and simplest implementation of a MiMe-NoCoDI loop. *G*_3_ corresponds to 1/*K*_*I*_ in the equation describing non-competitive inhibition. *x*: setpoint; *e*: control error; *u*: manipulated variable; *y*: controlled variable; *y*_*R*_ instantaneous value.

Combining the equations for the single blocks and solving for a selected signal delivers e.g. for the controlled variable *y*(*t*)

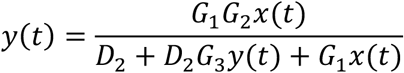

and for the manipulated variable *u*(*t*)

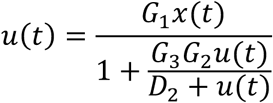

as recursive solutions for the temporal development of the system [3].

In order to arrive at an equifinal solution describing the steady-state behaviour of the systems (in terms of input and constant parameters only), we define

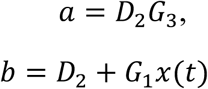

and

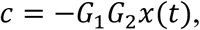

so that the equation for *y* may be expressed as

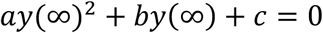

with the two solutions

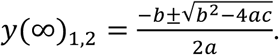

The positive solution represents the fixpoint (set point) of the feedback control system [3].

For *k* parallel feedback loops we obtain the recursive universal equation

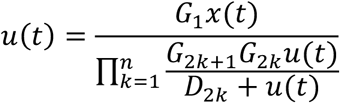

with the equifinal solution

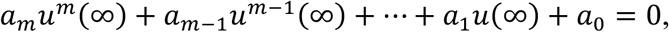

where *m* = *k* + 1 [3].

As an extension of this mathematical formulation, a class library was developed for the programming languages Object Pascal and S that facilitates the development of simulation programs for this kind of information processing structure. It has been made available as an open-source project (CyberUnits Bricks) [10]. The class hierarchy is shown in Fig. 2. Non-visual classes support numeric simulation, visual classes the generation of block diagrams.

**Figure 2.**
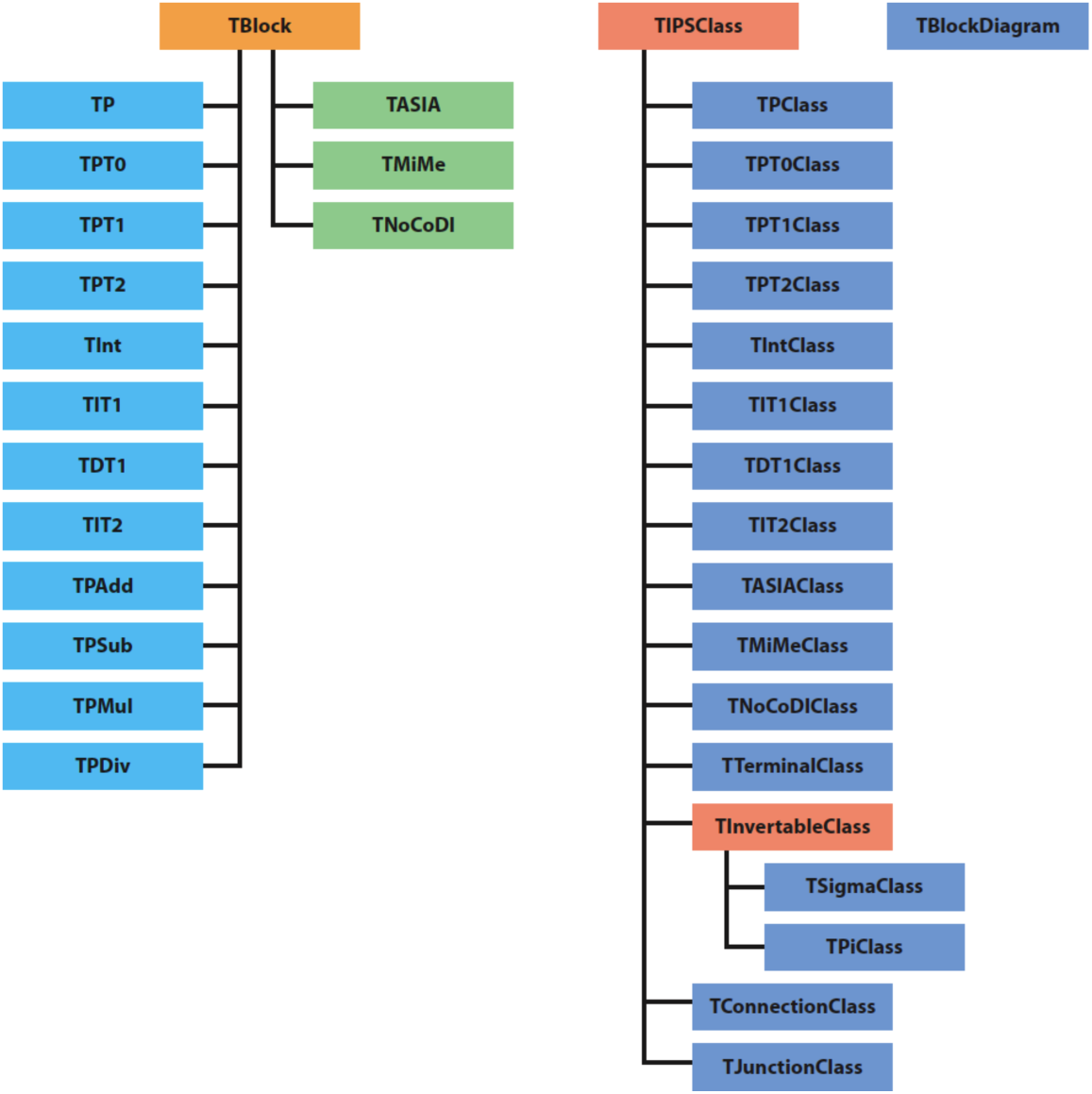
Class hierarchy of the CyberUnits Bricks library. Non-visual classes are shown on the left (descendants of *TBlock*), visual classes on the right (descendants of *TIPSClass*).

## RESULTS AND DISCUSSION

Parameters representing extensive properties can be derived from body size. If determined with biochemical research they can be translated to whole-body values with appropriate scaling factors, representing e.g. the mass or volume of organs or compartments. Examples include the gain factors (*G*) of the presented equations and the *α* parameters of ASIA elements.

The process of vertical translation is more complex for intensive properties, e.g. dissociation constants (*D* and *K*_*I*_ coefficients of the specified equations). It is important to note that *D* is a lumped parameter that incorporates the rate constants for the reactions involved in enzyme kinetics:

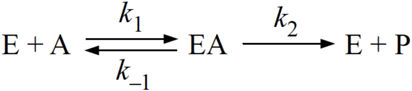

with

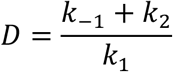

This can be used for translating from molecular properties to the whole-body behaviour of a feedback loop. In the example of the simple MiMe-NoCoDI loop shown in Fig. 1, this can be achieved by redefining a and b as

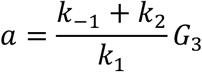

and

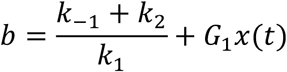

and using these parameters for the above-mentioned quadratic equation. Fig. 3 shows how the controlled variable depends on selected rate constants.

**Figure 3.**
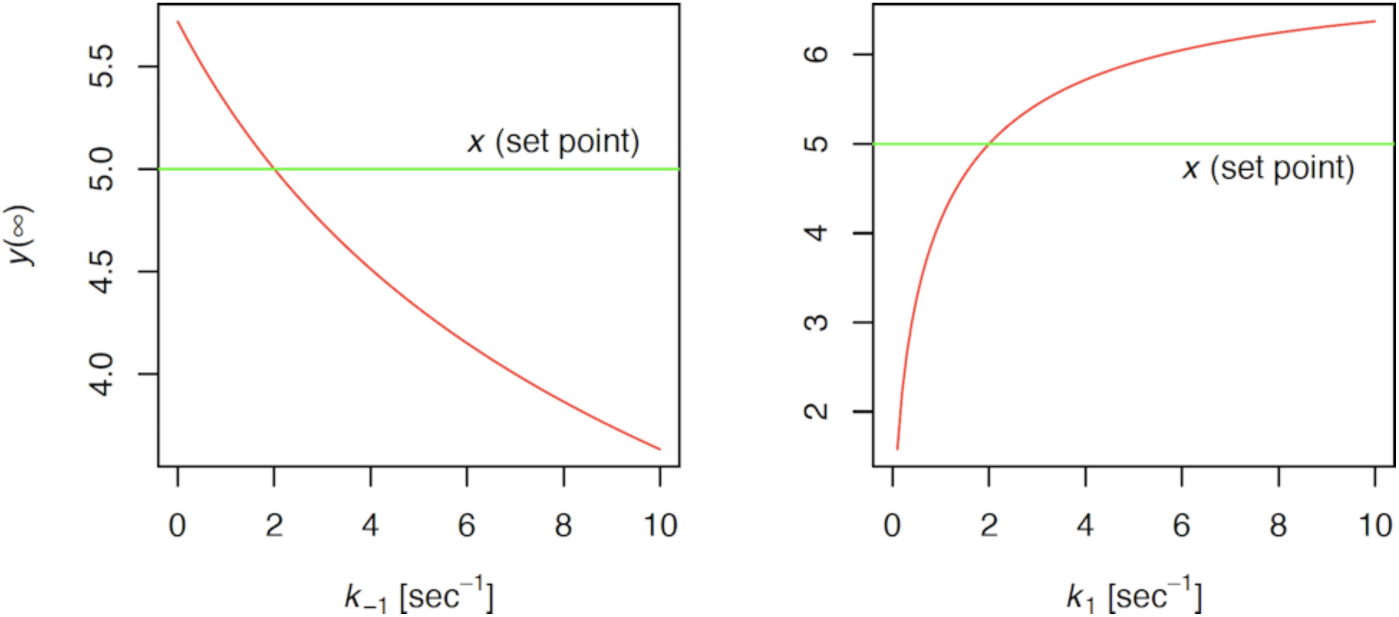
Controlled variable *y* in relation to the rate constants for the dissociation of the enzyme-substrate complex.

After appropriate rescaling, the same relationships also apply to statistical properties on the level of single molecules, since the rate constants can be expressed as the probability of the corresponding equation occurring in the subsequent time interval or as the inverse mean lifetime of the complex. By translating properties of a feedback loop to rate coefficients it is possible to verify the conclusions with computer simulations of molecular reactions, e.g. based on the Gillespie algorithm [11, 12], and to integrate this theory with the methodology of stoichiometric network analysis [13].

Some of the results shown here are still partly preliminary and require future research for a more systematic investigation and validation of the vertical relationships between causal levels.

If this method can be successfully implemented it might be a valuable tool, e.g. for the research on endocrine disruptors and genetic syndromes and for the early phases of drug design, which is more and more supported by computational methods.

## CONCLUSION

The method presented here allows for translating between molecular processes and the behaviour of embedding feedback loops on the level of the whole organism, a previously unsolved challenge. This is accomplished by “high-fidelity” modelling of feedback control systems based on a parametrically isomorphic approach and by applying the insights to the statistical properties of chemical reactions. The new methodology may be a useful tool for multiple fields of biomedical research.

## Acknowledgements

JWD is co-owner of the intellectual property rights for the patent “System and Method for Deriving Parameters for Homeostatic Feedback Control of an Individual” (Singapore Institute for Clinical Sciences, Biomedical Sciences Institutes, Application Number 201208940-5, WIPO number WO/2014/088516). The author wants to express his gratitude to Ljiljana Kolar-Anić (Faculty of Physical Chemistry, University of Belgrade) and Željko D Čupić (Department of Catalysis and Chemical Engineering, University of Belgrade) for valuable discussions and suggestions.

## Notes

http://cyberunits.sf.net

http://quantum-salis.sf.net

### Competing Interest Statement

The authors have declared no competing interest.

